# Geneticin shows selective antiviral activity against SARS-CoV-2 by interfering with programmed -1 ribosomal frameshifting

**DOI:** 10.1101/2022.03.08.483429

**Authors:** Carmine Varricchio, Gregory Mathez, Trestan Pillonel, Claire Bertelli, Laurent Kaiser, Caroline Tapparel, Andrea Brancale, Valeria Cagno

## Abstract

SARS-CoV-2 is currently causing an unprecedented pandemic. While vaccines are massively deployed, we still lack effective large-scale antiviral therapies. In the quest for antivirals targeting conserved structures, we focused on molecules able to bind viral RNA secondary structures. Aminoglycosides are a class of antibiotics known to interact with the ribosomal RNA of both prokaryotes and eukaryotes and have previously been shown to exert antiviral activities by interacting with viral RNA. Here we show that the aminoglycoside geneticin is endowed with antiviral activity against all tested variants of SARS-CoV-2, in different cell lines and in a respiratory tissue model at non-toxic concentrations. The mechanism of action is an early inhibition of RNA replication and protein expression related to a decrease in the efficiency of the -1 programmed ribosomal frameshift (PRF) signal of SARS-CoV-2. Using in silico modelling, we have identified a potential binding site of geneticin in the pseudoknot of frameshift RNA motif. Moreover, we have selected, through virtual screening, additional RNA binding compounds, interacting with the same site with increased potency.

## INTRODUCTION

Since the beginning of the SARS-CoV-2 pandemic, a huge effort has been made for the identification of effective vaccines and antivirals. The vaccines programme has been an immense success with the approval of three vaccines in less than one year, and the vaccination, at the time of writing, of 66.9% of the world population. (Statistic and Research Coronavirus Vaccinations, https://ourworldindata.org/covid-vaccinations, accessed July 2022). The drug discovery effort has also led to the identification of three antiviral drugs, Remdesivir, Molnupiravir and Paxlovid, which have been approved by the FDA (Parums, 2022). However, the emergence of new SARS-COV 2 variants, which can potentially escape the vaccine-mediated immunity and the effectiveness of therapies, highlights the importance to identify new potential pan antiviral agents against SARS-CoV-2.

RNA structure elements represent an attractive target for antiviral drug discovery. Viral genomes contain highly conserved RNA elements that play a critical role in gene regulation and viral replication. These RNA elements are directly involved in the viral infection process, interacting with proteins, DNA or other RNAs, modulating their activity (Embarc-Buh et al., 2021). The function and activity of these RNA molecules are based on the complex three-dimensional structure they can adopt (Ganser et al., 2019). Due to its conserved nature and its well-defined structure, the RNA provides potentially unique interaction sites for selective small-molecule ligands that affect viral replication. The high conservation of untranslated regions reduces the possibility of a drug-resistant mechanism, increasing the effectiveness of potential antiviral drugs (Warner et al., 2018). Any change in nucleotide sequence can result in inactive elements through misfolding the RNA structure, as recently demonstrated with the programmed -1 ribosomal frameshifting element (−1 PRF) of SARS-CoV-2 (Bhatt et al., 2021). Programmed ribosomal frameshifting is one of the strategies commonly used by RNA viruses, such as flaviviruses, coronaviruses, influenza A viruses and HIV, to regulate the relative expression level of two proteins encoded on the same messenger RNA (mRNA) (Brierley and dos Ramos, 2006; Firth et al., 2012; Penn et al., 2020). This strategy is rarely used by human cells, making it an attractive therapeutic target for antiviral drug development. Several studies have proposed the frameshifting element (FSE) as a target for disruption of virus replication (Ahn et al., 2011; Haniff et al., 2020; Park et al., 2011; de Wit et al., 2016). The SARS-CoV-2 FSE is a small region between the open reading frame (ORF) 1a and the ORF 1b. The ORF1b encodes all the enzymes necessary for viral RNA replication, including the RNA-dependent RNA polymerase. The frameshifting events depend on the flexibility of the RNA structure and its ability to interact with the ribosome. A small molecule that can alter the structural organisation of the FSE can block the frameshifting event and consequently the viral replication.

Aminoglycosides are among the molecules known to interact with secondary or tertiary structures on RNA, therefore potentially inhibiting -1 PRF of SARS-CoV-2. This class of antibiotics is known to interact with the ribosomal RNA of prokaryotes and eukaryotes (Garreau De Loubresse et al., 2014; Vicens and Westhof, 2003a) in particular with the tRNA recognition site, blocking a conformational switch of the ribosomal A site. The affinity for RNA makes this class of molecules potentially interact with additional RNA structures as shown for RNA HIV dimerisation sites, or for a riboswitch sequence in the 5’ leader RNA of a resistance gene in bacteria (Jia et al., 2013).

Among the different aminoglycosides, geneticin is one of the few for which the cells are permeable, and it is commonly used in cell lines as a selective agent due to its alteration in eukaryotic protein synthesis when administered at high doses for a prolonged time (Davies and Jimenez, 1980). However, the drug proved to be effective as well against multiple viruses (Bovine Viral Diarrhea Virus, Dengue Virus and Hepatitis C virus (HCV)) (Alexander V Birk1 et al., 2008; Ariza-Mateos et al., 2016; Zhang et al., 2009) at non-toxic concentrations. In particular, in the evaluation of the antiviral activity of geneticin against HCV, a specific interaction with a double-stranded RNA switch structure in the 5’UTR of the virus was shown (Ariza-Mateos et al., 2016), its binding resulted in a stabilisation of the open conformation leading to inhibition of the production of non-structural protein 3 (NS3) and viral replication in cell lines.

Here we show that geneticin is active against SARS-CoV-2 through an early inhibition in its life cycle and an alteration of the -1 PRF efficiency. The activity in the micromolar range is maintained against multiple variants, in different cell lines, and in respiratory tissues and has a high barrier to resistance. Importantly, we identified a putative binding site for geneticin on the -1 PRF sequence of SARS-CoV-2 through *in silico* modelling. After a screening of RNA binding molecules interacting with the same site, we identified compounds displaying antiviral activity at lower half-maximal effective concentrations (EC_50_) than geneticin, paving the road for the future development of SARS-CoV-2 antivirals.

## RESULTS

### Geneticin is active against different variants of SARS-CoV-2 at non-toxic concentrations

Antiviral activity of geneticin against several variants of SARS-CoV-2 was assessed in Vero-E6 cells with the addition of the molecule post-infection. Merafloxacin, a molecule previously shown to inhibit SARS-CoV-2 (Hoffmann et al., 2020), was tested against B.1.1.7 as control. Importantly the seven different variants tested, including the alpha (B.1.1.7), the beta (B.1.135), the delta (B.1.617.2) and the omicron (BA.1) were directly isolated from clinical specimens at the University Hospital of Lausanne with minimal passaging in cell lines to avoid any cell adaptation (Mathez and Cagno, 2021). We observed dose-response activity in the micromolar range for all the variants tested (Table 1). Analysis of the sequences did not reveal any particular cell adaptation, nor common changes in the variants showing higher EC_50_s if compared to the others (Supplementary Figure 1). Moreover, the activity of geneticin was conserved against HCoV-229e in Huh7 cells (Table 1), while we observed a lack of activity against an unrelated virus, namely Influenza A virus (H1N1) (Table 1). Importantly, we excluded that the antiviral activity is linked to a toxic effect of geneticin on the cell with viability, cytotoxicity, and apoptosis assays (Supplementary Figure 2A-B-C). Moreover, we verified that at the highest concentration used in the antiviral assays (600 µM) the protein synthesis in the cell is not impaired, in opposition with cycloheximide, a known elongation blocker (Supplementary Figure 2D).

**Table 1.** Antiviral activity of geneticin against SARS-CoV-2. EC_50_: half-maximal effective concentration. EC_90_: 90% effective concentration. 95% CI: confidence interval 95%

| | Virus | Variant | EC<sub>50</sub> (95% CI) [μM] | EC<sub>90</sub> (95% CI) [μM] | CC<sub>50</sub> [μM] |
| --- | --- | --- | --- | --- | --- |
| Geneticin | SARS-CoV-2 | B.1.1.7 (Alpha) | 47.2 (35.1 - 62.4) | 245 (149 - 460) | 3951 |
|  |  | B.1.351 (Beta) | 129 (84.7 - 188) | 542 (273 - 1537) | 3951 |
|  |  | B.1.617.2 (Delta) | 32.8 (21.3 - 48.2) | 201 (100 - 505) | 3951 |
|  |  | BA.1 (Omicron) | 25.7 (16.9 - 38.9) | 155 (60 - 528) | 934 |
|  |  | B.1.258 | 42.4 (29.7 - 60.4) | 163 (86 - 377) | 3951 |
|  |  | B.1.160 | 31.6 (21.2 - 44.8) | 150 (74 - 368) | 3951 |
|  |  | B.1.177 | 101.7 (74.8 - 138) | 407 (222 - 852) | 3951 |
|  | HCoV-229e |  | 7.21 (5.25 - 9.58) | 23.9 (12.1 - 62.3) | 684 |
|  | Influenza virus |  | > 600 |  | 5841 |
| Merafloxacin | SARS-CoV-2 | B.1.1.7 (Alpha) | 23.7 (16.8 - 33.8) | 116 (52.9 - 458) | > 100 |

### The antiviral activity is maintained in human respiratory cell lines and in tissues

To assess the antiviral activity in more relevant cell models, we evaluated the antiviral activity in dose response of geneticin in Calu3 cells, a lung adenocarcinoma cell line, which was previously shown to mimic faithfully SARS-CoV-2 infection in respiratory cell line (Thi Nhu Thao et al., 2020). The results evidenced a sustained antiviral activity (EC_50_ 179.6 µM) also in this cellular model in absence of toxicity (Figure 1A). We then tested the activity in a pseudostratified model of the human respiratory tract (Mucilair, Epithelix). This tissue model is composed of the typical cells of the human upper respiratory tract, namely ciliated, goblet and basal cells. In this infection model, we aimed to mimic a possible treatment with the molecule by starting the treatment 24 hours post-infection (hpi) when the infection of the tissue was already well established and we used viral stocks produced in the same tissue and never passaged in cell lines to exclude any adaptation. The treatment was performed apically and the infection was monitored up to 4 days post-infection by collecting an apical wash and performing either a qPCR or a titration in Vero E6 (Figure 1B). The results evidenced significant protection from viral infections with both B.1.1 (B) and omicron BA.1 (C) variants, without a decrease in viability for the tissues (Supplementary Figure 2).

**Figure 1.**
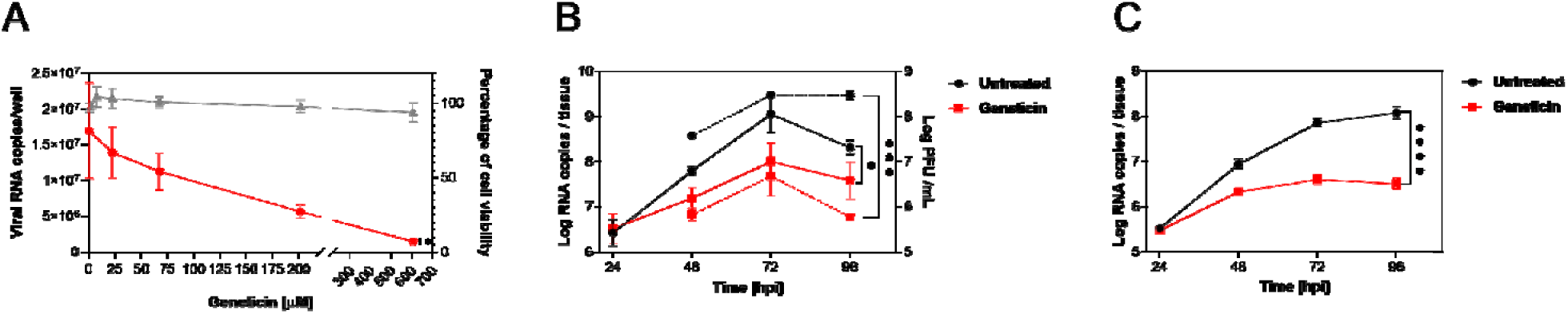
The activity of geneticin in maintained in Calu3 cells and in human-derived tissues. A) Calu3 cells were infected with SARS-CoV-2 (alpha variant) for 1 hour at 37°C. After the removal of the inoculum, the cells were treated with serial dilutions of geneticin. At 24 hpi supernatant was collected and viral RNA copies were evaluated with qPCR. B) and C) Mucilair tissues were infected with B) SARS-CoV-2 B.1.1 10^6^ RNA copies or C) SARS-CoV-2 Omicron BA.1 10^5^ RNA copies, the following day the apical treatment with 30 µg/tissue started. Every 24 hours an apical wash was performed and collected after 20 minutes at 37°C. The supernatant was then used for viral RNA quantification (solid lines) or for plaque assay (dashed lines). The results are the mean and SEM of two to three independent experiments performed in duplicate. P values <0.0332 (*), <0.0021 (**), <0.0002 (***), < 0.0001 (****)

### Geneticin has a high barrier to resistance

To evaluate the barrier for resistance, we passaged the virus in presence or absence of increasing doses of geneticin for 11 passages (Supplementary Figure 4A). At the end of the experiment, we evaluated the EC_50_s of the viruses grown in presence of geneticin in comparison to the untreated viruses. We did not observe any significant change in the EC_50_s. We verified through NGS the presence of mutations in the untreated and geneticin treated viruses. We observed the typical features of viruses passaged in cell lines such as the inactivation or deletion of the furin cleavage site (R685H in the untreated, del 679-85 in the geneticin treated), and we could identify a mutation in the ORF1a in the geneticin treated not present in the untreated at passage 11. However, the same mutation was not present in the duplicate condition (Supplementary Figure 4B).

### Geneticin inhibits the -1PRF of SARS-CoV-2

In order to assess the mechanism of action and the stage of viral replication of SARS-CoV-2 inhibited by geneticin, we first assessed viral protein expression. We exploited a GFP expressing SARS-CoV-2 previously generated (Thi Nhu Thao et al., 2020) evaluating the GFP expression in presence or absence of the drug at 24h and 48hpi (Supplementary Figure 5). The results evidenced, as expected, a marked reduction in the number of infected cells and in addition, the GFP intensity was significantly reduced in the infected cells treated with geneticin, when compared to the untreated control (Supplementary Figure 5). Moreover, we analysed the amount of viral nucleoprotein and cellular tubulin in cells infected and treated for 24 or 48h by western blot, confirming the marked selective reduction in viral protein expression (Figure 3A and Supplementary Figure 6). We included as control merafloxacin, a compound previously shown to interfere with the -1 PRF signal. The results show a decrease in viral protein production for both compounds, suggesting a block of infection at an initial stage of the viral life cycle. To evaluate if the inhibition of protein expression was related to a block of translation or viral replication, we then monitored viral replication through a RT-qPCR measuring the viral RNA replication at different time points. As shown in Figure 3B, the addition of geneticin or merafloxacin, results in inhibition of viral RNA replication at 4h, 8h and 24h post-infection, demonstrating a rapid inhibition of viral replication by the two drugs. Finally, we assessed the ability of geneticin, in comparison with merafloxacin, to interfere with the programmed ribosomal frameshifting element of SARS-CoV-2 with a dual luciferase assay. The -1 PRF signal was cloned between Renilla and Firefly luciferase and the relative expression of the luciferases was evaluated in presence or absence of the drugs as described in (Bhatt et al.). The results of figure 3C show a reduction in the -1 PRF efficiency in presence of both compounds suggesting a direct interaction of the drug with the - 1PRF sequence resulting in impaired replication (Figure 3B) and protein production (Figure 3A). Additionally, we tested mutated PRF as described in Bhatt et al. (Figure 4D). The results showed a similar inhibitory profile for geneticin and merafloxacin, suggesting a common binding pocket on the -1 PRF.

**Figure 2.**
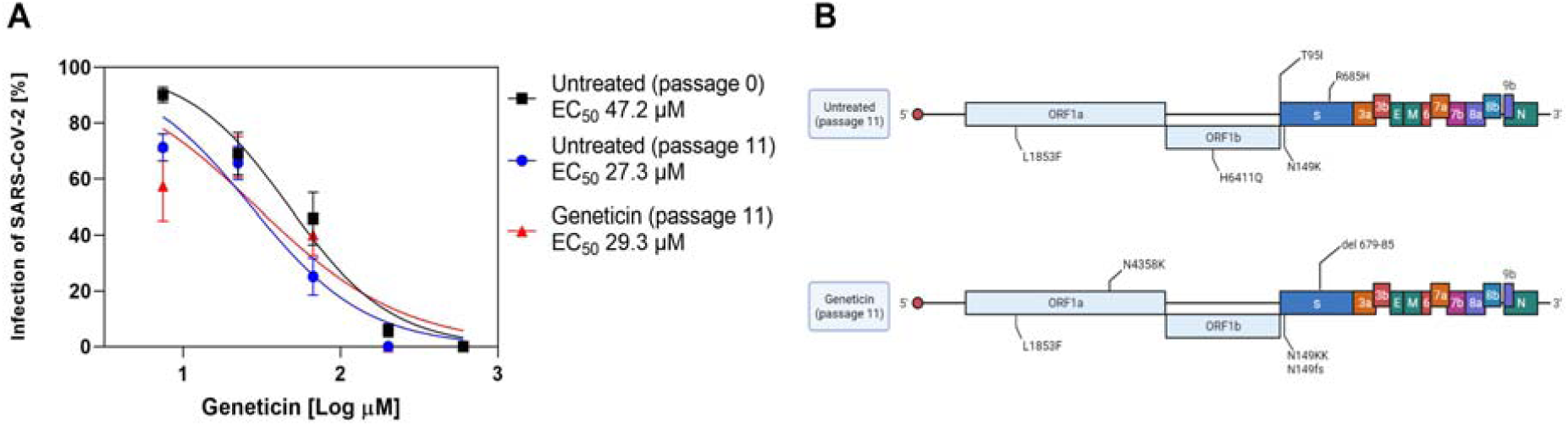
A) The EC_50_s of geneticin were evaluated against the B.1.1.7 stock, viruses are grown in Vero E6 without treatment for 11 passages, or in presence of increasing doses of geneticin. B) The mutations observed at passage 11 as compared to the original B.1.1.7 stock (created with biorender.com).

**Figure 3.**
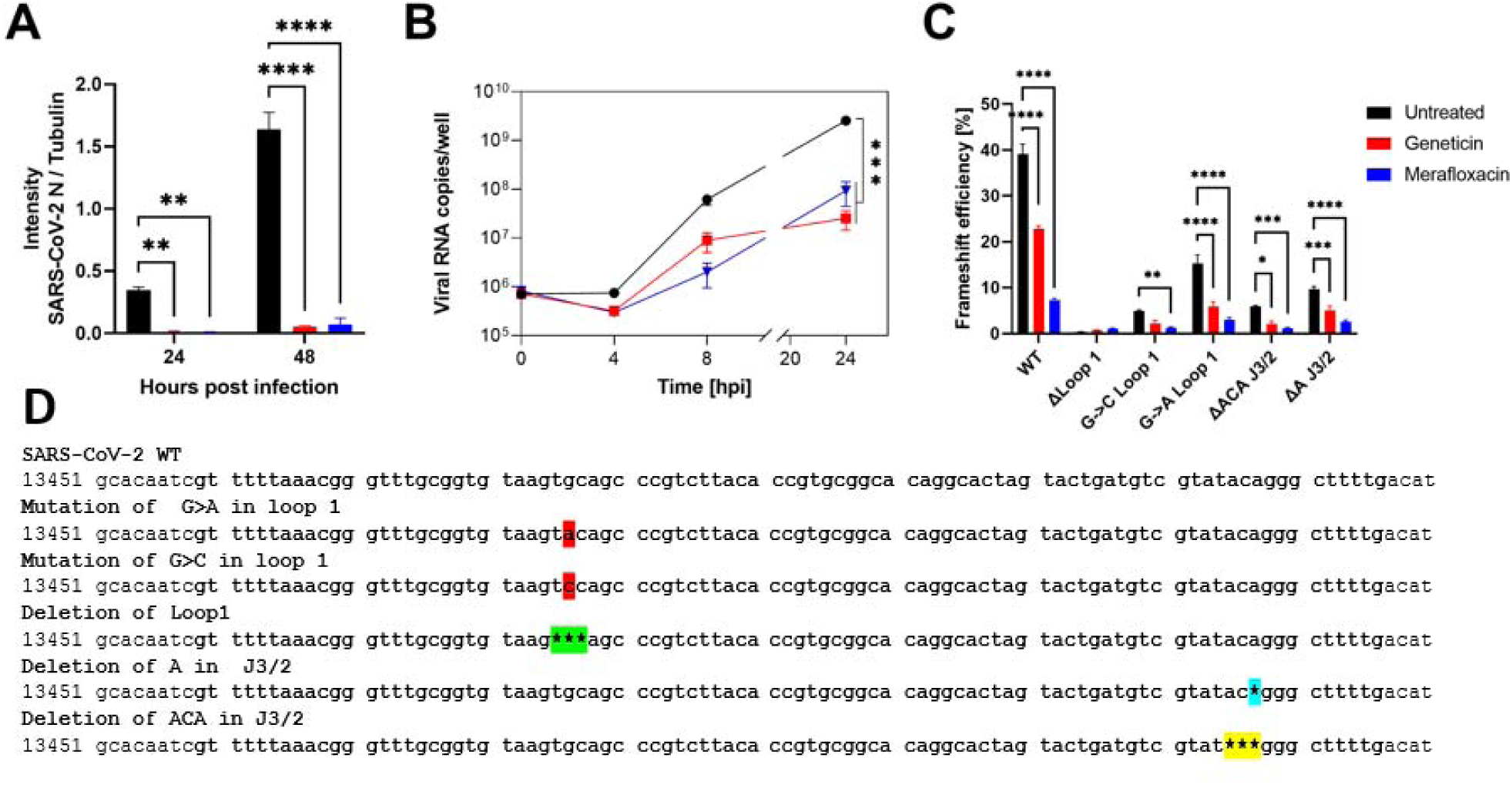
Mechanism of action of geneticin. A) Vero-E6 cells were infected with B.1.1.7 SARS-CoV-2 at MOI 0.1 and at MOI 0.01 (24hpi and 48hpi conditions respectively). Cells were treated post-infection with geneticin (600 µM). Cells were lysed 24hpi or 48hpi and protein quantification was done by Western Blot (Supplementary Figure 6). Values are expressed by the ratio of the intensity of SARS-CoV-2 nucleocapsid over alpha tubulin quantified by ImageJ. B) Vero-E6 were infected with SARS-CoV-2 at MOI 0.1 for 1 hour at 37°C. After the removal of the inoculum, geneticin (600 µM) or merafloxacin (100 µM) were added to the well. At 0, 4, 8 and 24 hours post.-infection cells were lysed and viral RNA was quantified. C) Dual luciferase evaluation was performed at 24 hours post-transfection in Vero-E6 cells treated with geneticin (600 µM) or merafloxacin (50 µM). The results are mean and SEM of three independent experiments performed in duplicate. P values <0.0332 (*), <0.0021 (**), <0.0002 (***), < 0.0001 (****). D) SARS-CoV-2 RNA frameshift-stimulatory element sequence and the mutant sequences.

**Figure 4.**
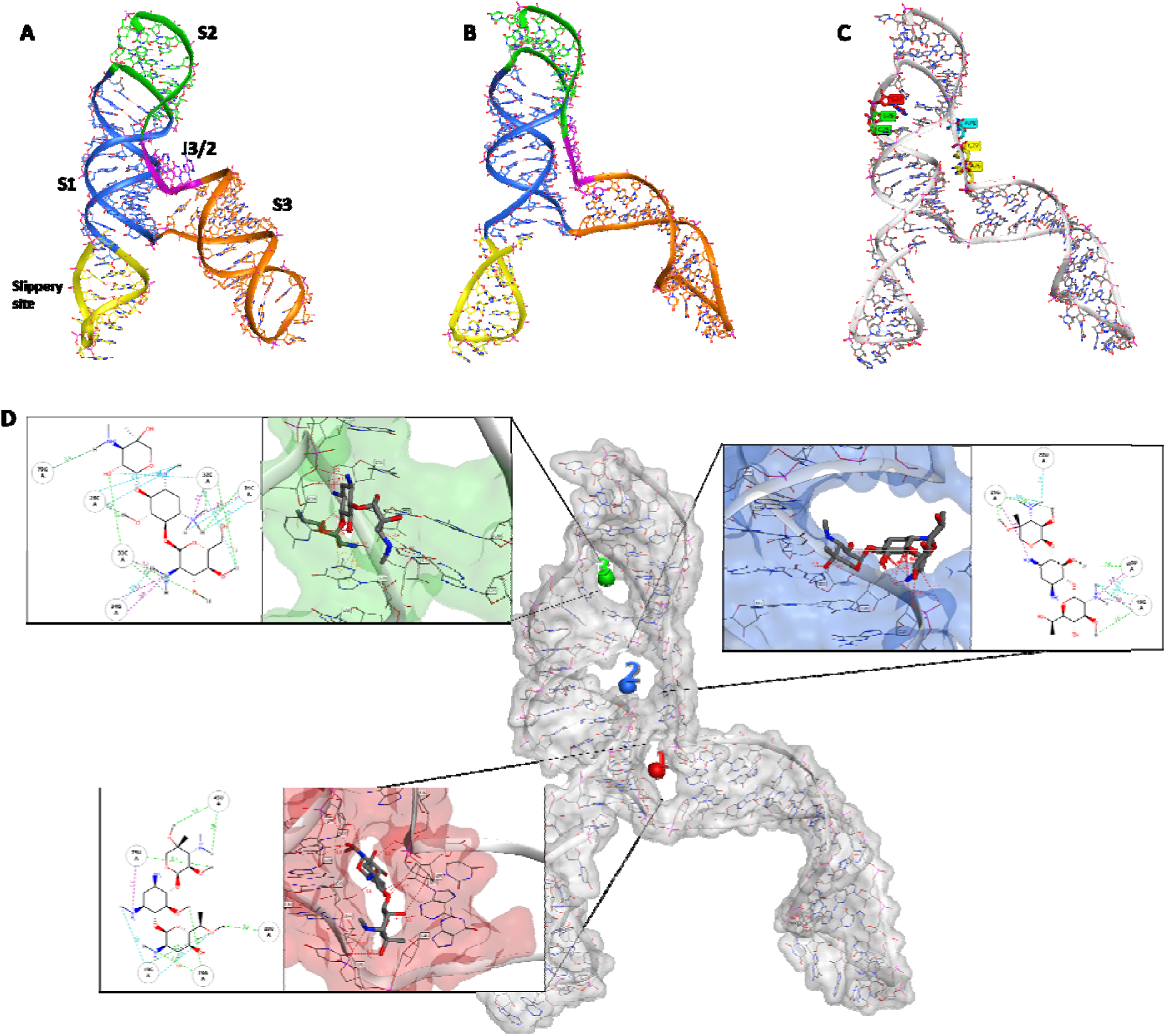
Comparison of the cryo-EM RNA structure (A) and the refined RNA structure by molecular dynamic simulation (B). (C) Different sequential mutations in the FSE structure: point mutations in Loop1 (in red); deletion of loop1 (in green); deletion of A and ACA in J3/2 (turquoise and yellow respectively). (D) The 3 binding sites identified by RNAsite. The binding site 1 (ring site); 2 (J3/2) and 3 (stem 2) are highlighted in red, blue and green, respectively. The 2D figures show geneticin interactions with the surrounding nucleotides in the binding sites. Turquoise dashed lines indicate weak H-bond; green dashed lines indicate strong H-bond, and purple dashed lines indicate electrostatic interactions

### *In silico* modelling and prediction of geneticin binding site

To further rationalise the results obtained by dual-luciferase and antiviral assays, the cryo-EM structure of the RNA frameshift-stimulatory element (FSE) was used to investigate the Geneticin-FSE binding complex (Zhang et al., 2021). The cryo-EM RNA structure shows a λ-like tertiary arrangement composed of a three-stemmed H-type pseudoknot structure with three loops. Starting from the 5’-end and proceeding to the 3’-end, the cryo-EM structure begins with a slippery site, followed by Stem 1 (S1), which leads to the Loop (L1), and it continues to Stem 2 (S2) (Figure 4A). From the second stem (S2), the RNA strands continue to form a hairpin region (S3), followed by an unpaired segment J3/2, which leads back to Stem 2 and closes the Stem 1-Stem 2 pseudoknot (Figure 4A). The cryo-EM data also suggested alternative conformations due to the structural flexibility at the 5’-ends, which appeared poorly resolved (Zhang et al., 2021). Moreover, the cryo-EM structure was resolved at low-mid resolution 6.9 Å, which can affect the assignment of the atom position with high certainty. Molecular dynamic (MD) simulations have proven useful in refining macromolecular structures, particularly unveiling the atomic details for low-resolution regions of the cryo-EM map (Bissaro et al., 2020; McGreevy et al., 2016; Nierzwicki and Palermo, 2021). In this study, we initially refined the cryo-EM FSE structure by 100 ns molecular dynamics simulation using the GROMACS software package (Abraham et al., 2015). Overall, after an initial 40 ns of equilibration, the structural fluctuation of the RNA reduced, with the simulation system converging around a fixed RMSD value of 1.5 Å. This RMSD value was chosen as a cut-off for selecting a series of different conformers, which were successively clustered to select a representative structure (Figure 4B). The comparison between the cryo-EM and our model showed a similar structure rearrangement with minimum RMSD variations in nucleotide position, except for the slippery site and S3 region, which displayed a higher level of flexibility (Figure 4B). These results are in line with previous studies conducted by Omar et al. and Rangan et al., showing that stem 3 could adopt multiple conformations (Omar et al., 2021a; Rangan et al., 2021).

Mutational studies showed that the virus replication is highly sensitive to any conformational change in the pseudoknots region, as evident by the point mutation of guanidine to adenine in loop 1 (Figure 3D and Figure 4C), which reduced the frameshifting efficiency to 60%. (Figure 3C). According to the mutation results and the uncertainty of the S3 region, we hypothesised that geneticin could significantly alter and disrupt the FSE conformational plasticity and consequently the viral replication, directly binding the S1/S2 -J3/2 pseudoknots region. A previous study showed that geneticin can interact with tertiary RNA structures through hydrogen bonds and electrostatic interactions (Prokhorova et al., 2017; Vicens and Westhof, 2003b). The binding affinity of geneticin for the RNA structures is mainly due to the presence of four amino groups which are positively charged at physiological pH and can form strong electrostatic interaction with the negatively charged phosphates in the nucleic acid backbone (Figure 4D). Furthermore, the presence of seven hydroxyl groups can stabilise the RNA-binding complex through a series of hydrogen bonds with the base atoms and phosphate oxygen atoms of the nucleic acid. Several studies demonstrated the preference of aminoglycoside compounds to bind RNA helix and junction sites (Aradi et al., 2020). We first investigated if there were potential geneticin-binding sites in FSE regions using the refined cryo-EM structure. The binding site analyses, performed by the RNAsite module (Su et al., 2021a), identified 3 different potential active sites situated between stem 1, stem 2 and junction site (Figure 4D), which partially confirmed the results obtained by Zhang and collaborators, who reported the presence of a ‘ring site’, a ‘J3/2 site’ and the ‘slippery hairpin binding site (Zhang et al., 2021). Our results showed that two potential binding sites, 1 and 2; which were located in close proximity, sharing 3 nucleotide residues (G18, G19 and G20), similar to the ring site and J3/2 site reported by Zhang and collaborators (Zhang et al., 2021). However, contrary to Zhang and collaborators, we could not detect any suitable binding site on the slippery site, instead, a new potential pocket (binding site 3) was located at the beginning of stem 2 (Figure 4D).

The geneticin-binding affinity was evaluated against all the three potential binding sites using an *in silico* protocol, which comprises three steps: firstly, the compound was docked using XP GLIDE module (Maestro, Schrodinger), then the docked poses were refined using MM-GBSA module, and lastly the refined poses were rescored using two scoring functions optimised specifically for RNA-ligand complex, Annapurna and Amber score function (DOCK6). The purpose of multiple scoring functions was to ascertain the most potentially accurate ligand poses and avoid any possible bias associated with using a single docking program/scoring function. The docking results showed that although geneticin can be well accommodated inside all three binding sites in different rational configurations, it has a slighter higher affinity for site 1 compared to sites 2 and 3 (ΔG_mm-gbsa_ -102.98, - 90.34, -80.77 kcal/mol, respectively). Site 2 and 3 showed the largest surface area, but are solvent-exposed, which affect the ligand-RNA interaction: geneticin was only partially in contact with the RNA surface while the rest of the molecule was exposed to solvent (Figure 4). On the other hand, site 1 showed a smaller surface area, but it was surrounded by nucleotides (G18, G19, G20, G43, G44, G46, U75 and A76), which form a tunnel-like binding site. Geneticin can well occupy the active site with the streptamine core inside the tunnel cavity, interacting with G19, U20, U45 and A74 through hydrogen bonds (H-bonds) and with U75 by electrostatic interactions between the amino group chain and phosphate groups of U75 (Figure 4D). To confirm the results obtained by the MM-GBSA analysis, the refined docked poses were rescored using Annapurna and DOCK6 score function. In both software, the top-ranked binding poses were predicted to site 1 (Supplementary Table 1), suggesting that this site might be more accessible and druggable than the other two binding sites. Interestingly, merafloxacin can also bind site 1, showing similar binding interactions of Geneticin (H-bonds with U45 and A74 and electrostatic interaction with U75) (Supplementary Figure 7A). The deletion of ACA nucleotides in the J2/3 regions (Figure 4C) is in close proximity to binding site 1, but it is not directly involved in the binding. Although it could potentially affect the RNA folding (supplementary Figure 7B), the dual luciferase assay did not show any decrease in the efficiency of merafloxacin nor geneticin in presence of this deletion (Figure 3C).

### Identification of -1PRF binding compounds

To test the druggability of the binding site, we screened an RNA-targeted library (Enamine, ChemDIV), which contains 44520 commercially available RNA-binding compounds, against site 1. The virtual screening was performed using the previously described protocol. Firstly, the XP glide docking mode was employed to virtually screen the RNA-target library. The best 10% of docked poses to this initial screening were refined and rescored through MM-GBSA. To validate the top-scored docking results, the compounds were rescored using Annapurna and DOCK6 scoring functions. After applying a consensus score procedure, 132 molecules were chosen, which were further evaluated by visual inspection considering the ability of compounds to occupy the binding site and the number of interactions formed between the compounds and the target. At the end of this workflow, twenty compounds were selected, purchased and evaluated in antiviral assays. Among them, three compounds could inhibit the virus replication with an EC_50_ in the micromolar range, with higher potency than geneticin (Figure 5A).

**Figure 5.**
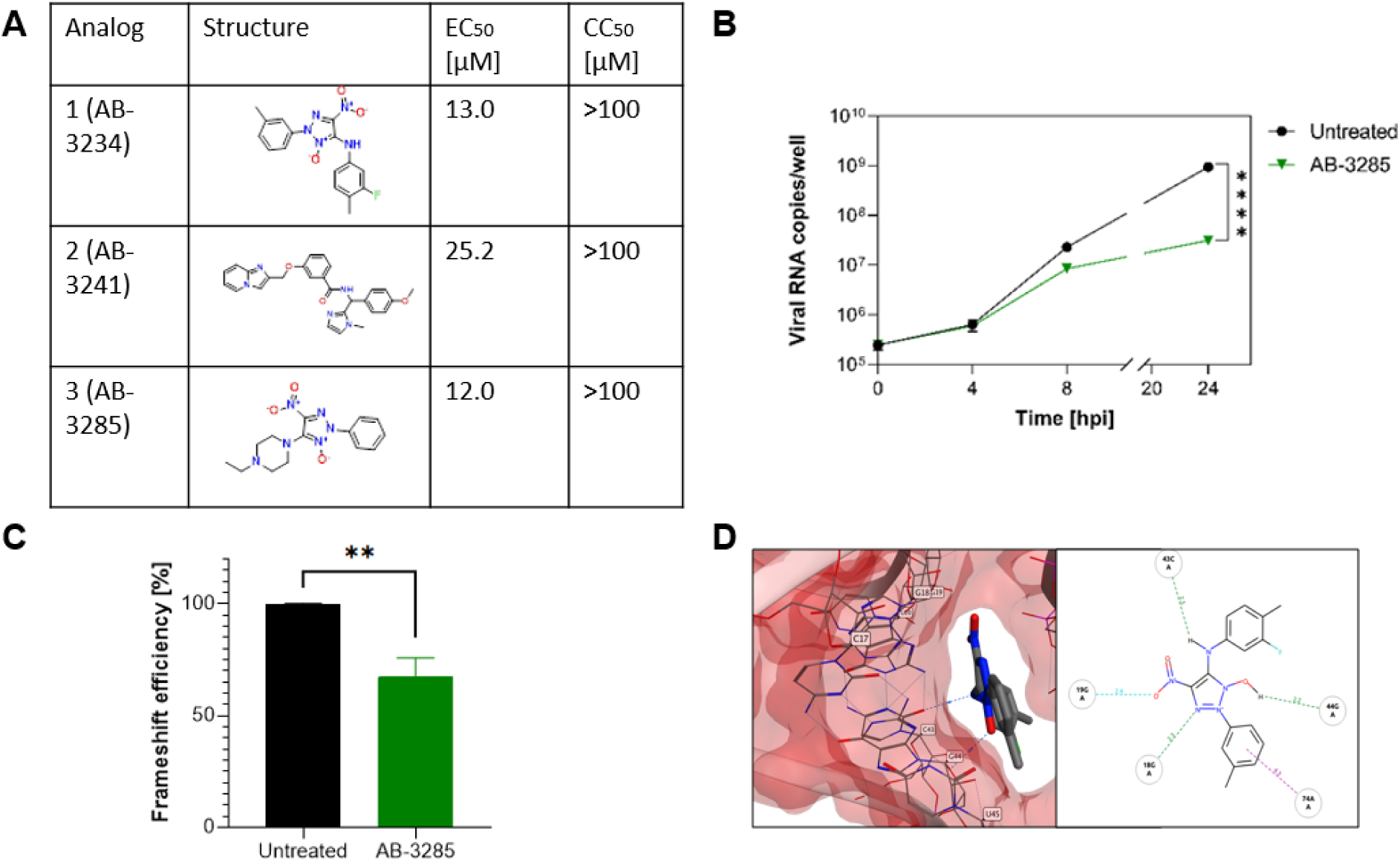
Antiviral activity of site 1 -1 PRF binders against SARS-CoV-2. A) Table of analogs with increased potency against B.1.1.7 SARS-CoV-2 than geneticin. EC_50_: half-maximal effective concentration, CC_50_: half-maximal cytotoxic concentration. B,C,D) Mechanism of action of AB-3285. B) Vero-E6 were infected with SARS-CoV-2 at MOI 0.1 for 1 hour at 37°C. After the removal of the inoculum, AB-3285 (250 μM) was added to the well. At 0, 4, 8 and 24 hours post.-infection cells were lysed and viral RNA was quantified. C) Dual luciferase evaluation was performed at 24 hours post-transfection in Vero-E6 cells treated with AB-3285 (500 µM). The frameshift efficiency was normalized compared to untreated. D) Binding pose of AB-3285 in the PRF binding site 1. The results are mean and SEM of at least two independent experiments performed in duplicate. P values <0.0332 (*), <0.0021 (**), <0.0002 (***), < 0.0001 (****)

The most potent compound was further analysed for kinetics of RNA expression (Figure 5B) and dual luciferase (Figure 5C) confirming a similar activity to geneticin. The *in-silico* results showed that could completely occupy the tunnel-binding site, forming a cation-pi with G19 and H-bonds with G19, G18, C43 and G44 (Figure 5D).

More recently, the SARS-COV-2 FSE structure solved by x-ray confirmed the cryo-EM three-stemmed H-type pseudoknot structure, but it showed different tertiary arrangements: the cryo-EM structure has a λ-like tertiary arrangement, meanwhile, the x-ray adopts a vertical conformation (Roman et al., 2021) Although the x-ray shows a higher resolution of 2.09 Å, it lacks the 5’-slippery site sequence, which might affect the tertiary arrangement. These different arrangements of the FSE have also been supported by previous chemical probing, mutational, and NMR studies demonstrating that the arrangement of stem 1 and stem 2 relative to stem 3 can be flexible (Schlick et al., 2021a). The superposition of the x-ray structure and our model showed a similar binding site in the experimental structure, as also revealed by RNAsite, which overlaps our identified binding site 1 (Supplementary Figure 8).

## DISCUSSION

The alteration of the flexibility of the FSE of SARS-CoV-2 is detrimental to the replication of the virus (Bhatt et al.; Huston et al., 2021; Manfredonia et al., 2020; Omar et al., 2021b; Schlick et al., 2021b). If the viral RNA cannot interact correctly with the ribosome, the -1 PRF is altered and ORF 1ab cannot be expressed at the correct ratio, resulting in a lack of production of the viral polymerase and a consequent reduction of the replication (Bhatt et al.). The FSE of SARS-CoV-2 was previously shown to be a possible target for antiviral development with basic modelling (Park et al., 2011) or with empiric screening with dual luciferase assays (Sun et al., 2020). The precise druggable pockets of the FSE however were not previously identified. With the aim of identifying new molecules interacting with viral RNA, we tested geneticin, an aminoglycoside known to interact with RNA secondary structures.

The compound proved to be effective against multiple variants of SARS-CoV-2 (Table 1). The range of EC_50_s determined (Table 1) might be linked to the fitness of the variants in the Vero E6 and their plaque-forming ability. To exclude any bias, we verified as well the activity of geneticin in Calu3 cells, and we tested both B.1.1 and omicron BA.1 variants in a human respiratory airway model. In all conditions, we confirmed the antiviral activity of geneticin (Figure 1). Furthermore, the compound showed activity as well against HCoV-229e, demonstrating a broad-spectrum activity against coronaviruses, while it was not active against an unrelated RNA virus, Influenza A virus (Table 1), proving that the mechanism of action is not related to a general effect on the ribosome that will impair the replication of all viruses. The absence of toxicity at the antiviral tested doses is demonstrated as well by the toxicity analysis (Figure S1) and by the western blot analysis in which we observe a significant decrease of viral nucleoprotein and an absence of effect on cellular tubulin (Figure 3 and Supplementary Figure 6).

We then verified an early inhibition in the life cycle with reduced viral protein expression and RNA replication (Figure 3A-B), and we tested the activity on the PRF through dual luciferase assays (Figure 3C) on WT and mutated sequences, in which geneticin and merafloxacin behaved similarly supporting the same mechanism of action and possibly the same binding site. Targeting a highly conserved sequence in the RNA, the development of resistance is intrinsically limited, however, we verified it by growing the virus in presence of increasing concentrations of geneticin; after 11 passages we failed to observe any difference in the EC_50_s nor the appearance of specific mutations, confirming the high barrier to resistance (Figure 2).

The high flexibility and plasticity of the FSE is an essential requirement for its biological activity (Bhatt et al.; Huston et al., 2021; Manfredonia et al., 2020; Omar et al., 2021b; Schlick et al., 2021b). This unique characteristic is also supported by cryo-EM and x-ray structures recently published (Bhatt et al.; Roman et al., 2021; Zhang et al., 2021). In particular, the pseudoknot structure seems to be highly dynamic before encountering the ribosome. However, the unique 3-stem architecture of the FSE (Figure 4) and its mechanism made the FSE a viable target for small molecules. Our computational studies confirmed the presence of a suitable binding site in the pseudoknot structure, originally identified by Zhang and collaborators (Zhang et al., 2021). This binding site is located between J3/2 and stem 3 regions, and it is large enough to accommodate geneticin, merafloxacin, and small ligands. Interestingly, this pocket is close to the S3 region of the FSE. Our molecular dynamic simulation studies revealed that the S3 region is particularly flexible, showing higher fluctuations than the other regions. According to these results, we hypothesised that the S3 region might play a critical role in the conformational change of the FSE, necessary for the frameshifting event. Hence, geneticin could exert its antiviral activity by altering the flexibility of this region, and consequently interfering with the conformational changes between the two main FSE structures.

The resulting antiviral activity is however linked to several limitations: the activity is in the micromolar range, and further studies should focus on the identification of more potent compounds. The antiviral activity is at non-toxic concentrations, also in human-derived respiratory tissues (Supplementary Figure 2), however, the selectivity index of geneticin is narrow since it is known to bind eukaryotic ribosomes and it is associated with toxicity in cell culture. Although the administration in a viral infection is most likely to be for a short duration, future work should be directed to the identification of compounds devoid of interaction with ribosomal RNA. Moreover, aminoglycosides are associated with nephrotoxicity and ototoxicity when administered systemically, therefore a topical administration should be envisaged for compounds similar to geneticin.

For these reasons, and to validate the druggability of the binding pocket identified, we used a virtual screening simulation to identify additional molecules, from a library of RNA binders. Our *in silico* screening against the “J3/2- stem 3” site revealed that the architecture of the pocket might be sufficiently complex to be targeted by more specific ligands. Through our simulations, we have identified molecules that might engage the FSE targeting the J3/2- stem 3 pocket, enhancing or reducing the pseudoknot stability. The identification of compound 3 with increased potency, reduction of RNA replication, and alteration of the -1PRF (Figure 7) demonstrates the feasibility of our approach. Future work will be directed toward the identification of analogues with increased potency, retaining the same mechanism of action, but suitable pharmacological properties.

## SIGNIFICANCE

Programmed -1 ribosomal frameshifting plays major functional and regulatory roles in the SARS-CoV-2 replication. Thus, the frameshifting element is an attractive target for the development of new potential antiviral drugs. In this study, we have shown that geneticin, a well-known aminoglycoside antibiotic, could inhibit the viral replication engaging the frameshifting element, similarly to merafloxacin, a structurally different molecule, previously reported in literature. The mode of action was confirmed by three different biological assays: the inhibition of RNA synthesis, the reduction of viral protein production, and the luciferase expression assays under control of the -1PRF in presence of WT and mutated sequences. Moreover, we have hypothesized a potentially targetable pocket in the FSE structure, which can well accommodate the geneticin as evident by the high *in silico* binding affinity. The druggability of the binding pocket identified was confirmed through an *in silico* screening of a small library of RNA-binding small molecules. Among them, one compound showed higher antiviral activity and lower cytotoxicity than geneticin.

## STAR+METHODS

- Key resource table
- Resource availability
- Method details
- Statistical analyses

## AVAILABILITY

Raw data are available at 10.6084/m9.figshare.20338695

## ACKNOWLEDGEMENT

We thank the diagnostics of the Institute of Microbiology from the University Hospital of Lausanne for providing the clinical specimens of SARS-CoV-2.

## Key resources table

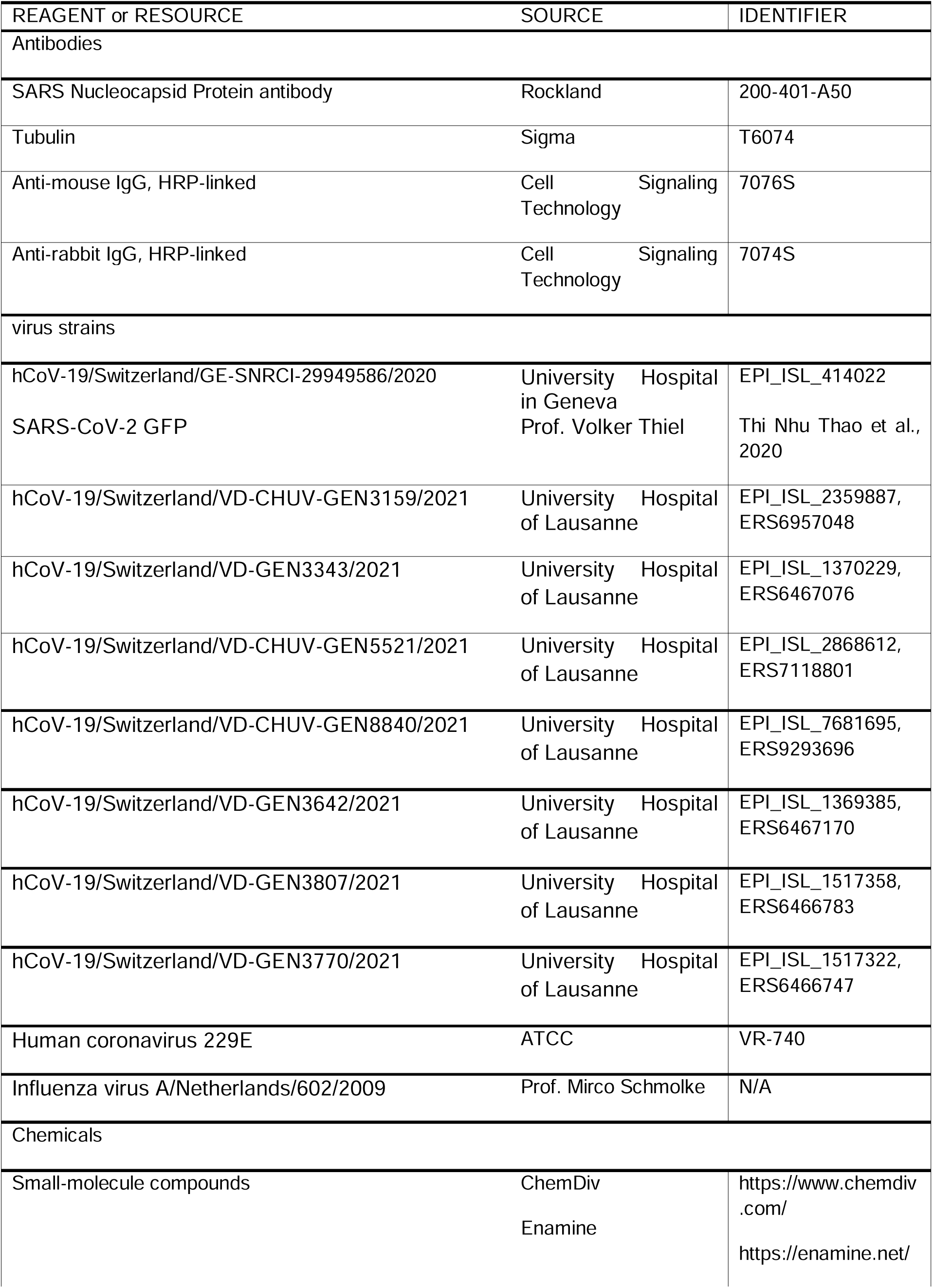
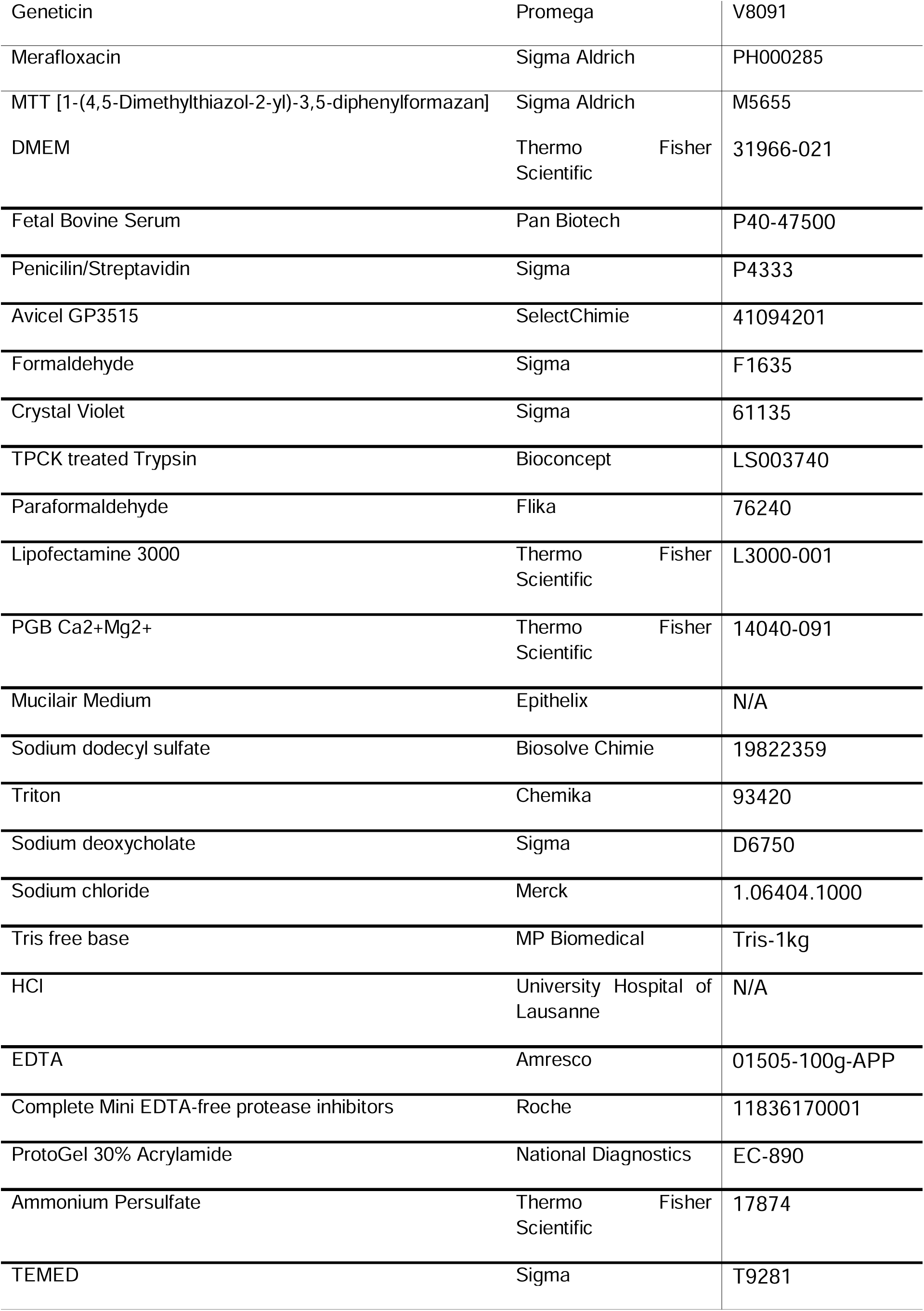
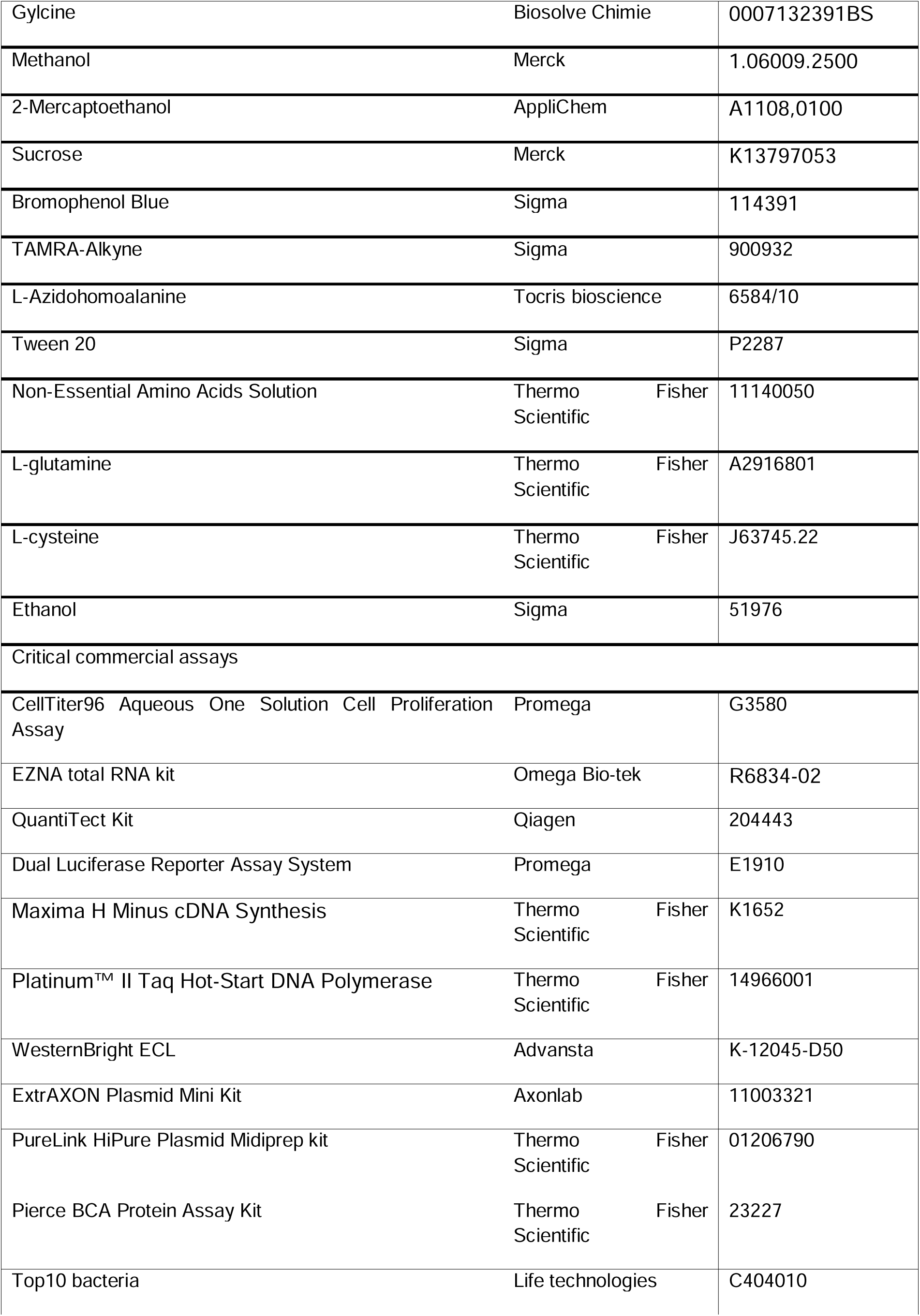
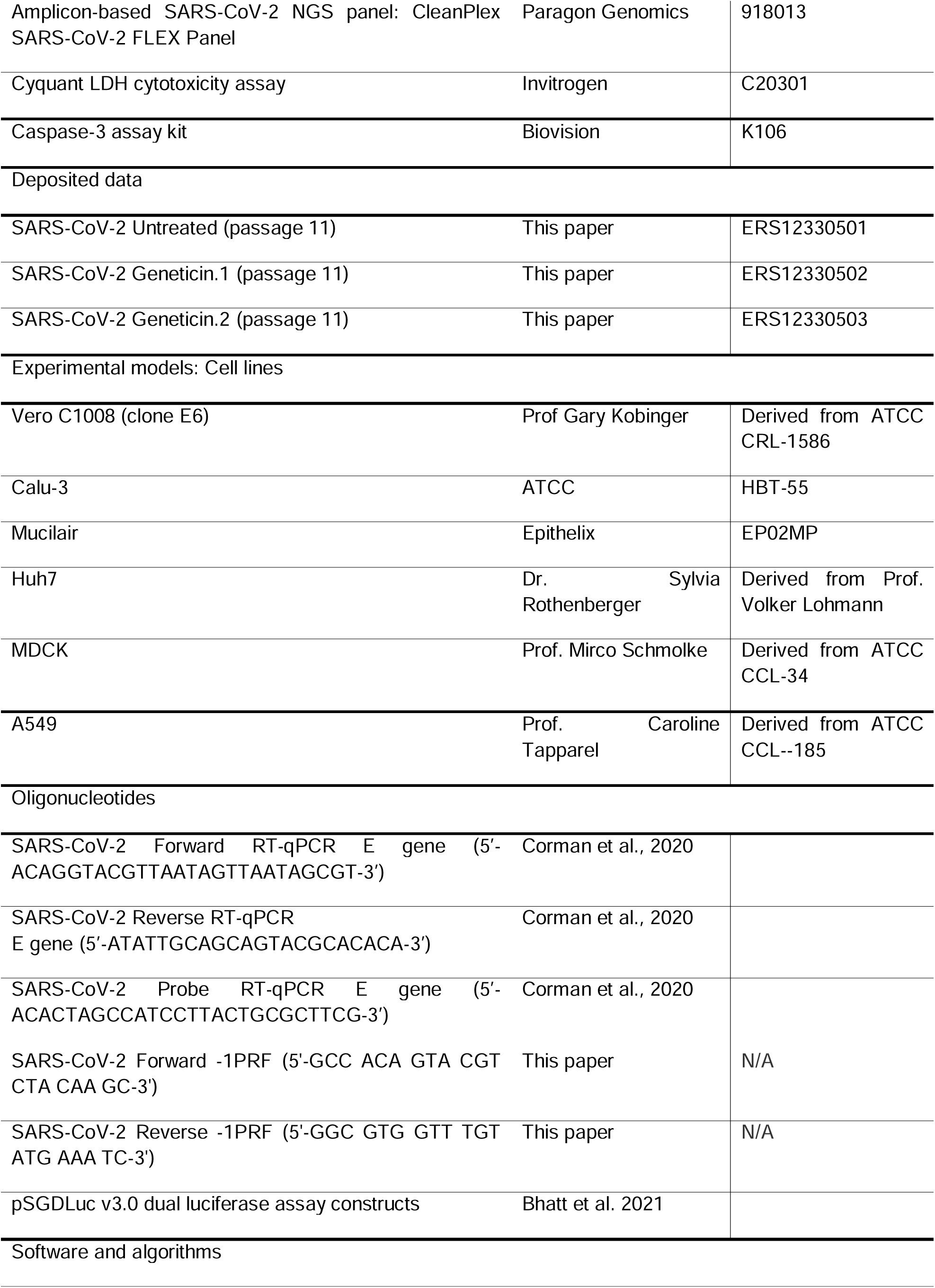
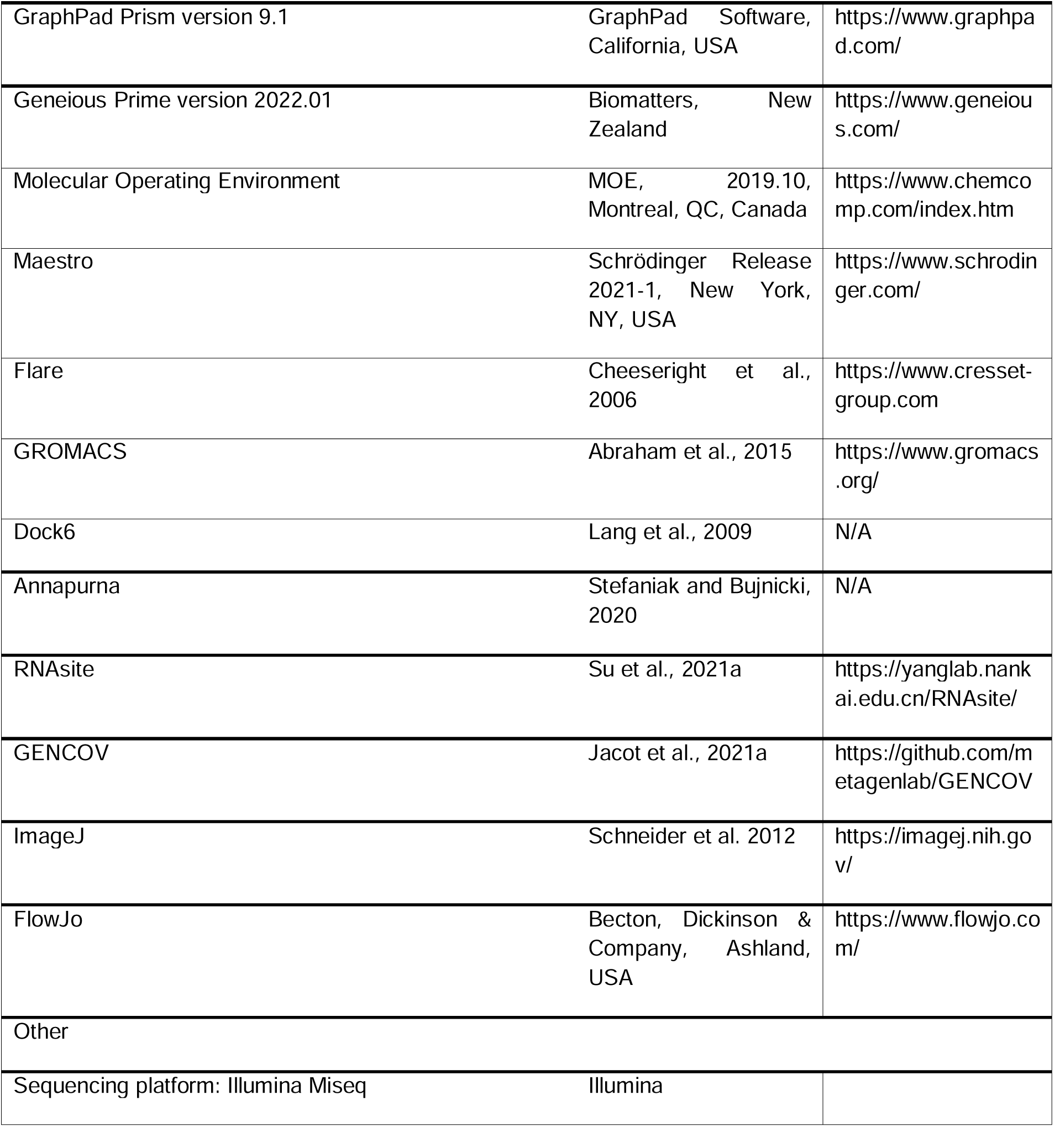

## MATERIAL AND METHODS

### Cells

Vero C1008 (clone E6) (ATCC CRL-1586), Huh7, MDCK and A549 cells were a kind gift from Prof Gary Kobinger, Dr Sylvia Rothenberger, Prof. Mirco Schmolke and Prof. Caroline Tapparel respectively. Calu-3 were purchased from ATCC. Cells were propagated in DMEM High Glucose + Glutamax supplemented with 10% fetal bovine serum (FBS) and 1% penicillin/streptavidin (pen/strep).

### Viruses

hCoV-19/Switzerland/GE-SNRCI-29949586/2020 (B.1.1) was isolated from a clinical specimen in the University Hospital in Geneva in Vero-E6 and passaged twice before the experiments. SARS-CoV-2 GFP was a kind gift from Prof Volker Thiel (Thi Nhu Thao et al., 2020). The other clinical strains (hCoV-19/Switzerland/VD-CHUV-GEN3159/2021 (B.1.1.7), hCoV-19/Switzerland/VD-GEN3343/2021 (B.1.351), hCoV-19/Switzerland/VD-CHUV-GEN5521/2021 (B.1.617.2), hCoV-19/Switzerland/VD-CHUV-GEN8840/2021 (BA.1), hCoV-19/Switzerland/VD-GEN3642/2021 (B.1.160), hCoV-19/Switzerland/VD-GEN3807/2021 (B.1.177), hCoV-19/Switzerland/VD-GEN3770/2021 (B.1.258)) were isolated from clinical specimens from the University Hospital of Lausanne (CHUV) as described in (Mathez and Cagno, 2021). The supernatant of infected cells was collected, clarified, aliquoted, and frozen at -80°C and subsequently titrated by plaque assay in Vero-E6.

### Cell toxicity assay

Cell viability was measured by the MTT assay or MTS assay (Promega) for tissues. Confluent cell cultures seeded in 96-well plates were incubated with different concentrations of geneticin in duplicate under the same experimental conditions described for the antiviral assays. Absorbance was measured using a Microplate Reader at 570 nm. The effect on cell viability at different concentrations of geneticin and additional compounds was expressed as a percentage, by comparing the absorbance of treated cells with the one of cells incubated with equal concentrations of solvent in medium. The 50 % cytotoxic concentrations (CC_50_) and 95 % confidence intervals (CIs) were determined using Prism software (Graph-Pad Software, San Diego, CA). For LDH assays the supernatants of the cells treated with geneticin as described before were analysed with cytotoxicity detection kit (CyQUANT™ LDH Cytotoxicity Assay, Thermofisher) and the 100% was calculated with a well in which the supernatant contained 0.05% triton. For apoptosis assay, the cells were analysed with the Caspase-3 assay kit (Biovision) according to the manufacturer instructions.

### Bioorthogonal Noncanonical Amino Acid Tagging (BONCAT)

A549 cells (350000 cells/well) were seeded in 6 well plate incubated with 600 µM of geneticin or 50 µg/ml of cycloheximide in medium without aminoacids supplemented with 1% NEAA, 1% glutamine, 50 µM L-cysteine and 50 µM L-Azidohomoalanine. Following 24h, 48h or 72h incubation wells were detached, pelleted fixed and subjected to click reaction with TAMRA-alkyne according to (Dieterich et al., 2007). The TAMRA signal was quantified with Cytoflex instrument and quantified with FlowJo.

### Antiviral assay in Vero-E6 cells

Vero-E6 cells (10^5^ cells per well) were seeded in 24-well plate. Cells were infected with SARS-CoV-2 (MOI, 0.001 PFU/cell) for 1 hour at 37°C. The monolayers were then washed and overlaid with medium supplemented with 5% FBS containing serial dilutions of compounds for the experiments with SARS-CoV-2 expressing GFP. For experiments with the different SARS-CoV-2 variants and analogues of geneticin, Vero-E6 cells were overlaid instead with 0.4% avicel gp3515 in medium containing 2.5% FBS. Two days after infection, cells were fixed with 4% formaldehyde and stained with crystal violet solution containing ethanol. Plaques were counted, and the percent inhibition of virus infectivity was determined by comparing the number of plaques in treated wells with the number in untreated control wells. 50% effective concentration (EC_50_) was calculated with Prism 9.1 (GraphPad).

### Antiviral assay in Calu3 cells

Calu-3 cells (4 × 10^4^ cells per well) were seeded in 96-well plate. Cells were infected with B.1.1.7 SARS-CoV-2 (MOI 0.1 PFU/cell) for 1 hour at 37°C. The monolayers were then washed and overlaid with medium containing serial dilutions of geneticin. At 24 hpi, supernatant was collected and viral RNA was extracted with EZNA total RNA kit (Omega Bio-tek). SARS-CoV-2 RNA was quantified by RT-qPCR with the QuantiTect Kit (Qiagen, 204443) with Sarbeco E gene primers and probe in a QuantStudio 3 thermocycler (Applied Biosystems). Percent inhibition of virus infectivity was determined by comparing viral load in treated wells with the viral load in untreated control wells. EC_50_ was calculated with Prism 9.1 (GraphPad).

### Resistance selection and next generation sequencing

Vero-E6 cells (3.5 × 10^5^ per well) were seeded in 6-well plate. At the first passage, cells were infected with B.1.1.7 SARS-CoV-2 (MOI 0.01) for 1 hour at 37°C. The inoculum was removed and an overlay with 2.5% FBS in DMEM was added. Half of EC_50_ concentration of geneticin (40 µM) was added in 2 wells. Two other wells were left untreated. Supernatant was collected 3 days post-infection and clarified at 2×10^3^ rpm for 5 min. Each sample was quantified by plaque assay in Vero-E6 cells (10^5^ cells per well) with an overlay of 0.6% avicel gp3515 in 2.5% FBS DMEM. For the following passages, cells were infected with the previous corresponding passage (MOI 0.01). The concentration of geneticin was doubled up to a final concentration of 600 µM.

RT-PCR targeting the PRF sequence was done for each condition at passage 10 (see below). At passage 11, untreated and treated conditions were used for dose-response with geneticin as described above. A sample per condition was lysed with TRK Lysis Buffer (Omega Bio-tek) for next generation sequencing as previously described (Jacot et al., 2021b). Briefly, SARS-CoV-2 genome was amplified with the CleanPlex® SARS-CoV-2 FLEX panel. The tiled amplicons were then sequenced with 2×150 bp on a MiSeq instrument (Illumina, San Diego, USA). Reads were analyzed with GENCOV https://github.com/metagenlab/GENCOV), a pipeline modified from CoVpipe (https://gitlab.com/RKIBioinformaticsPipelines/ncov_minipipe). Variant calling was performed with Freebayes (Garrison and Marth, 2012) (parameters: –min-alternate-fraction 0.1 –min-coverage 10 – min-alternate-count 9) and consensus sequences were obtained using bcftools (Danecek et al., 2021) based on variants supported by at least 70% of reads. Lineages were assigned to the consensus sequence using Pangolin (O’Toole et al., 2021)

### Kinetics of RNA expression

Vero-E6 cells were seeded in 24-well plates at a density of 10^5^ cells per well and infected in duplicate with B.1.1.7 SARS-CoV-2 at MOI 0.1 PFU/cell for 1 hour at 37°C. After the removal of the inoculum the treatment was started and cells were lysed with TRK buffer (Omega Biotech) at 0, 4, 8 and 24 hours post infection. RNA was extracted with the Total RNA kit (Omega Biotech) and amplified with the E-sarbeco primers for SARS-CoV-2.

### Western Blot

Vero-E6 were seeded in 6-well plates at a density of 3.5*10^5^ cells per well. Cells were infected with B.1.1.7 SARS-CoV-2 with a different MOI for each time point (MOI 0.1 for 24hpi and MOI 0.01 for 48hpi). After one hour of infection at 37°C, inoculum was removed and fresh DMEM 2.5% FBS was added. Geneticin 600 µM and merafloxacin 50 µM were used as post-treatment. One or two days after infection, cells were lysed at 4°C for 30min with RIPA buffer (0.001% SDS, 0.01% Triton, 0.1% sodium deoxycholate, 5µM NaCl, 0.0025% Tris HCl, 2nM EDTA, protease inhibitors) and clarified at 13’000rpm for 30min. Supernantants were collected and quantified with Pierce BCA Protein Assay Kit (Thermo Fisher Scientific). Twenty µg of proteins were loaded in an 8% acrylamide gel (8% acrylamide, 0.05% SDS, 422mM Tris HCl, 0.1%APS, 0.001% TEMED) and separated at 150V for 2 hours in running buffer (0.1% SDS, 25mM Tris, 190mM glycine). Proteins were transferred on a nitrocellulose membrane after 1 hour at 100V in transfer buffer (20% methanol, 50mM Tris, 40mM glycine, 0.037% SDS). Nitrocellulose membrane was blocked with 5% milk diluted in TTBS (0.05% Tween, 20mM Tris HCl, 500mM NaCl) for 30min at room temperature. The membrane was incubated overnight at 4°C with 1:5000 alpha-tubulin and 1:5000 SARS-CoV-2 nucleocapsid antibodies in TTBS with 5% milk. After three washes in TTBS, 1:2000 anti-mouse IgG and 1:2000 anti-rabbit IgG HRP-linked antibodies was added on the membrane. The membrane were developed with WesternBright ECL (Advansta). Intensity of alpha tubulin and SARS-CoV-2 nucleocapsid were quantified by ImageJ.

### Flow cytometry analysis

Vero-E6 cells were seeded in 24-well plates at a density of 10^5^ cells per well and infected in duplicate with SARS-CoV-2 GFP at an MOI of 0.01 PFU/cell for 1 hour at 37°C. The cells were then treated with geneticin and incubated at 37 °C for additional 24 or 48 hours. Supernatant was collected, cells washed once and detached with trypsin. Once in suspension cells were pelleted and then fixed with paraformaldehyde 4% in PBS. Percentages of GFP positive cells and mean GFP value for each positive cell was evaluated with an Accuri C6 cytometer (BD biosciences).

### Dual luciferase

pSGDLuc v3.0 was modified to include the -1 PRF signal of SARS-CoV-2 as described in (Bhatt et al.). Vero-E6 cells were seeded 24 hours in advance in 96-well plates (104 cells per well), treated with geneticin, merafloxacin or geneticin analog and transfected with Lipofectamine 3000 (Thermofisher) and the plasmid containing the -1PRF sequence or the in frame control. Luciferase was evaluated 24 hours post transfection with the Dual Luciferase Reporter Assay System (Promega). The percentage of ribosomal frameshift was calculated as described in (Bhatt et al.).

### PRF sequencing

RNA was extracted from isolated clinical SARS-CoV-2 with E.Z.N.A total RNA (Omega Bio-Tek). Maxima H Minus cDNA Synthesis (Thermofisher) and Platinium II Taq (Thermofisher) were used as RT-PCR kits with designed primers (Fwd 5’-GCC ACA GTA CGT CTA CAA GC-3’, Rev 5’-GGC GTG GTT TGT ATG AAA TC-3’). PCR products were Sanger sequenced by Microsynth.

### MucilAir antiviral assays

Tissues were obtained from Epithelix (Geneva, Switzerland). For all experiments, epithelia were prepared with different single donor’s biopsies. Before inoculation with the viruses, MucilAir tissues were incubated in 250 μL of PBS Ca2+Mg2+ (PBS++) for 45 min at 37°C. Infection was done with 10^6^ RNA copies/tissue with B.1.1 SARS-CoV-2 or 10^5^ RNA copies with SARS-CoV-2 BA.1 (omicron). At 4 hours after incubation at 33°C, tissues were rinsed three times with MucilAir medium to remove non-adsorbed virus and cultures were continued in the air-liquid interface. Every 24 hours, 200 μL of MucilAir medium was applied to the apical face of the tissue for 20 min at 33°C for sample collection, followed by apical treatment with geneticin (30 µg/tissue) starting at 24 hpi. Viral load was determined by qPCR as described previously. At the same time point, the basal medium was replaced with 500 μL of fresh MucilAir medium. At the end of the experiments, tissues were fixed and subjected to immunofluorescence.

### Statistics and data analysis

Experiments were performed in duplicate and from two to four independent experiments as stated in the figure legends. Results are shown as mean and SEM. The EC_50_ and CC_50_ values for inhibition curves were calculated by regression analysis using the program GraphPad Prism version 9.1 to fit a variable slope sigmoidal dose-response curve as described in (Mathez and Cagno, 2021). One-way or Two-ways Anova followed by multiple comparison analysis was used as statistical tests to compare grouped analysis. Unpaired t test was used to compare two different conditions. Area under the curve analysis followed by unpaired t test or one-way ANOVA was done to compare curves.

### Molecular Modelling

All molecular modelling experiments were performed on Asus WS X299 PRO Intel® i9-10980XECPU @ 3.00GHz × 36 running Ubuntu 18.04 (graphic card: GeForce RTX 2080 Ti). Molecular Operating Environment (MOE, 2019.10, Montreal, QC, Canada); Maestro (Schrödinger Release 2021-1, New York, NY, USA); GROMACS (2020.4) (Abraham et al., 2015); Dock6 (Lang et al., 2009); Annapurna (Stefaniak and Bujnicki, 2020); RNAsite (Su et al., 2021a), were used as molecular modelling software. A library of commercially available RNA-targeting compounds was downloaded from Enamine and ChemDiv website.

### Molecular dynamic simulations

MD simulations were performed with Gromacs software package. The ff99+bsc0+χOL3 force field was used for MD simulation since this is the most validated and recommended FFs for RNA system (Aytenfisu et al., 2017). The cryo-EM of the SARS-COV-2 FSE was download from PDB (http://www.rcsb.org/; PDB entry 6xrz). The structure was solvated with 14,0812 TIP4P-Ew waters and 87 Na+ counterions to neutralise the charge on the RNA. All the molecular dynamics simulations were performed for 100ns on the isothermal-isobaric ensemble, using the stochastic velocity rescaling thermostat at 300 K and the Berendsen barostat with an isotropic pressure coupling of 1 bar. The RMSD, was used to verify the stability of the simulated systems during the MD simulation. The conformations obtained after 40 ns were extracted, and RMSD between all structures was used to perform a cluster analysis to group the different RNA conformations and to select a representative structure.

### Binding site identification and molecular docking

The refined cryo-EM structure was prepared for further refinement with the Schrödinger Protein Preparation Wizard. Protonation states of RNA nucleotides were calculated considering a temperature of 300 K and a pH of 7.4, and restrained energy minimisation of the added hydrogens using the OPLS4 force field was performed. The Geneticin and the RNA-targeting compounds were prepared using the Maestro LigPrep tool by energy minimising the structures (OPLS4 force field), generating possible ionisation states at pH 7 ± 2 (Epik), tautomers and stereoisomers per each ligand. RNAsite was employed to identify a potential binding site using the refined structure(Su et al., 2021b). An 11 Å docking grid was prepared using as the centroid the predicted binding pocket previously identified by RNAsite. A Glide XP precision was employed to screen the compounds keeping the default parameters and setting 3 as the number of output poses per input ligand. The best-docked poses were then refined using MM-GBSA module. The docking poses obtained were then rescored using Annapurna and amber DOCK6 scoring functions. The values of the three different scoring functions for each docking pose were then analysed together (consensus score) and only the Docking poses falling in the top 25% of the score value range in all the three scoring functions were selected for the final visual inspection. The visual inspection process, conducted as the last step of the structure-based virtual screening, was performed using MOE 2019.10. The 2D interaction plot was generated using Flare

