## Supplementary Data for "Geneticin shows selective antiviral activity against SARS-CoV-2 by interfering with programmed -1 ribosomal frameshifting"

Valeria Cagno

Institute of Microbiology of Lausanne

Rue du Bugnon 48

1011 Lausanne

+41213142611

† The authors wish it to be known that, in their opinion, the first 2 authors should be regarded as joint First Authors

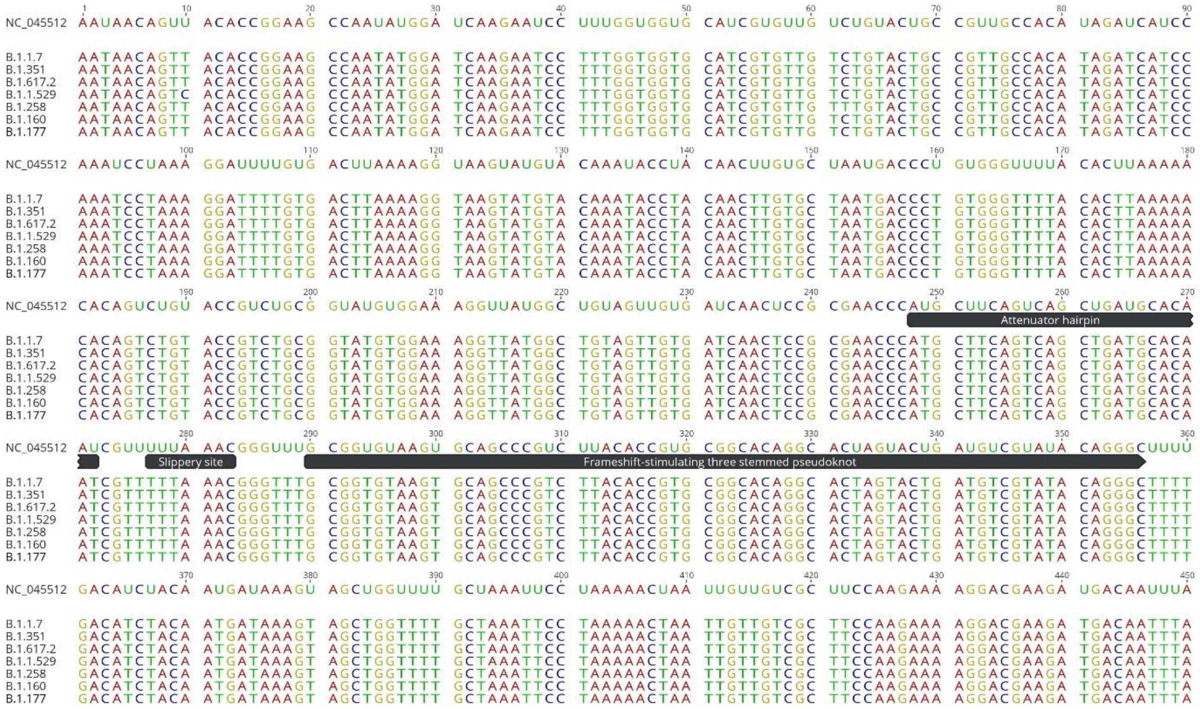


Supplementary Figure 1. Alignment of the different variants of SARS-CoV-2 with the reference SARS-CoV-2 isolated in Wuhan (NC_045512). The alignment shown correspond to nucleotides 13'186 to 13'635 of the reference.

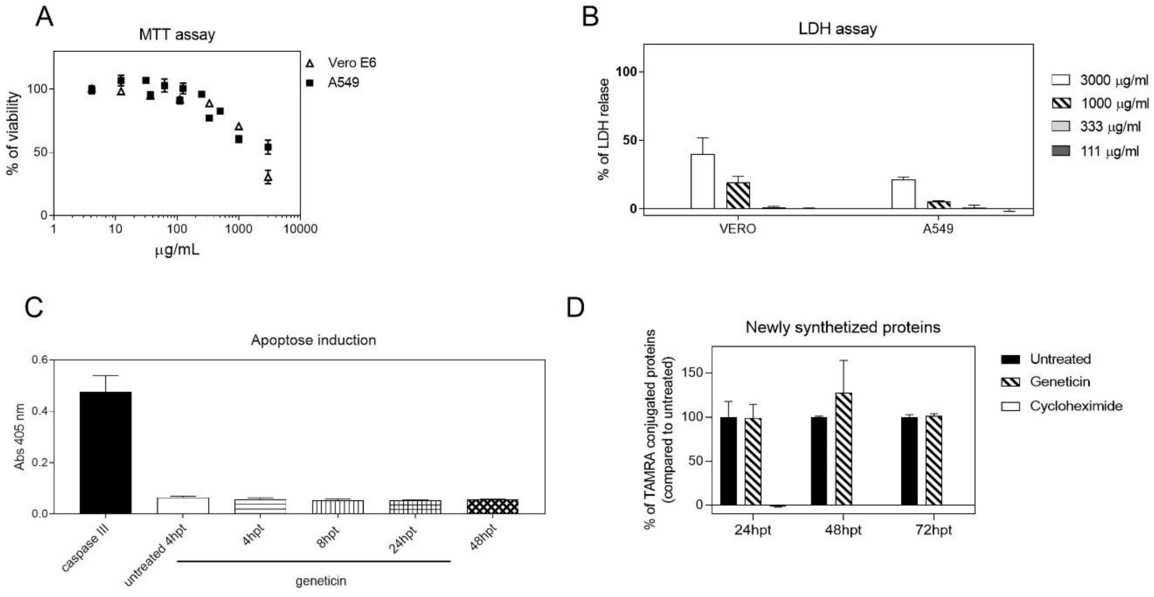


Supplementary Figure 2. Cell viability was evaluated with MTT assay (A), LDH release assay (B), Apoptose assay in Vero cells (C) and the amount of newly synthetized proteins was quantified with BONCAT in A549 cells treated with geneticin (600 µM) or cycloheximide (50 µg/ml) (D).

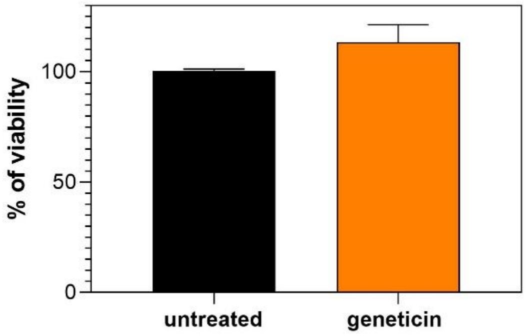


Supplementary Figure 3. Mucilair tissues were subjected to MTS assay at 96hpi according to manufacturer instructions. Results are mean and SEM of 2 independent experiments.

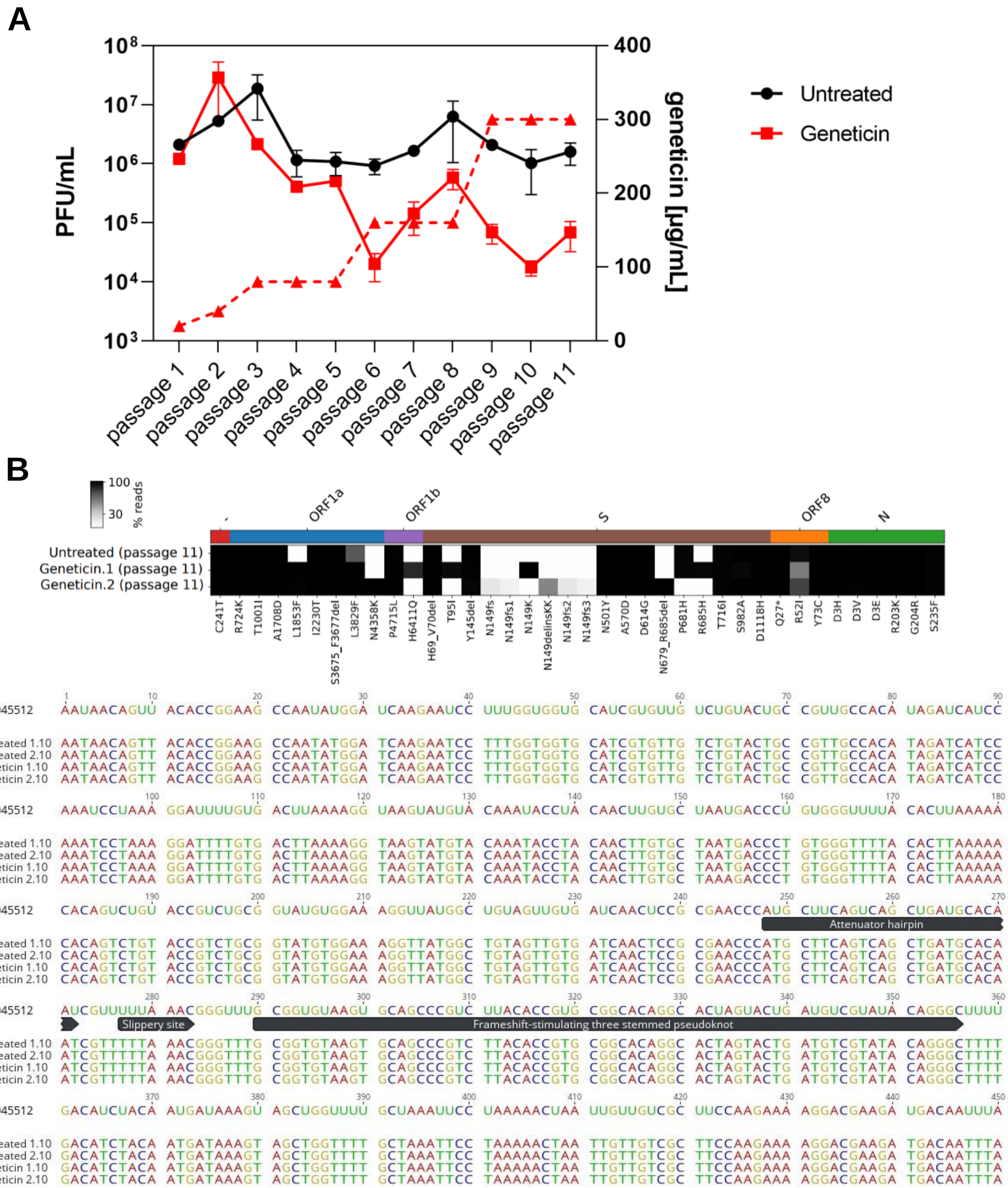


Supplementary Figure 4. Resistance against geneticin. A) Titration of untreated and geneticin treated passages (solid lines) with corresponding concentration of drug (dashed line). B) Mutations observed at passage 11 after next generation sequencing are represented in the top. Alignment of the different samples collected at passage 10 with the reference SARS-CoV-2 isolated in Wuhan (NC_045512). The alignment shown correspond to nucleotides 13’186 to 13’635 of the reference.

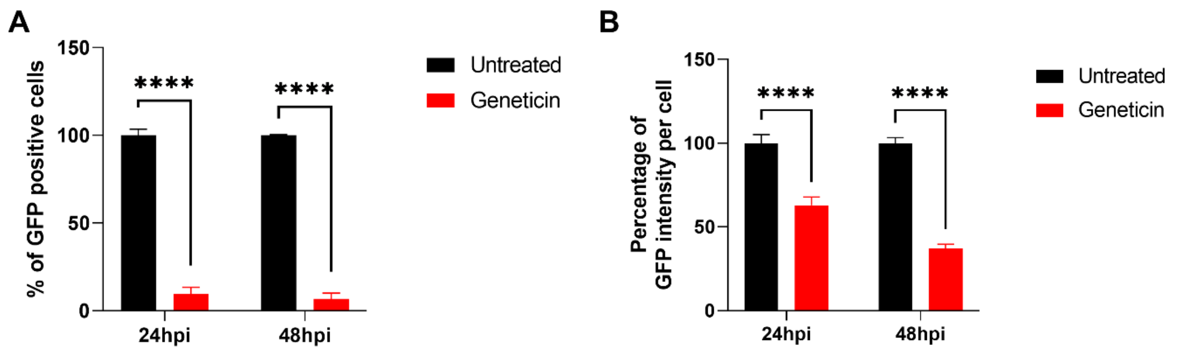


Supplementary Figure 5. SARS-CoV-2 GFP inhibition. VeroE6 cells were infected with SARS-CoV-2 expressing GFP at MOI 0.01, after the removal of the viral inoculum cells were treated with geneticin 300 µg/ml. At 24h and 48h cells were trypsinized and analyzed with a flow cytometer. Results are expressed as percentage of untreated control. ****P< 0.001

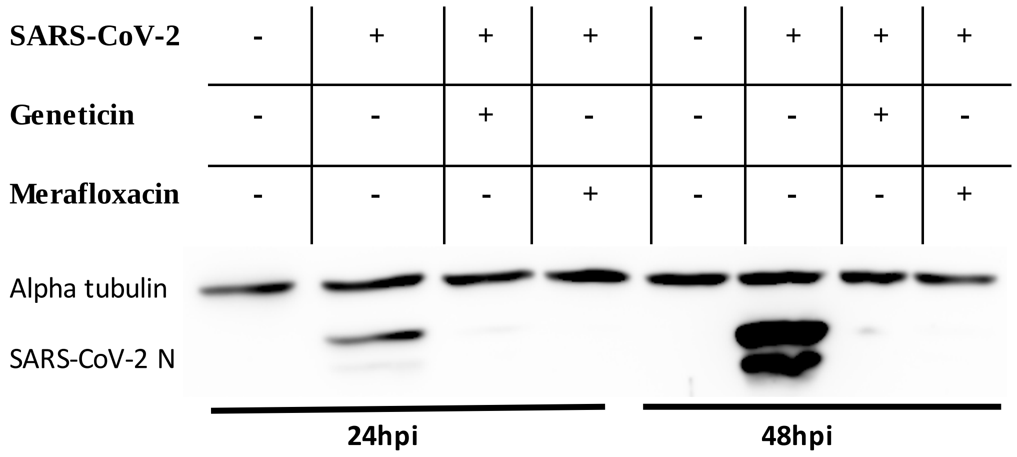


Supplementary Figure 6. Blockade of protein expression by geneticin. VeroE6 cells were infected with B.1.1.7 SARS-CoV-2 at MOI 0.1 and at MOI 0.01 (24hpi and 48hpi condition respectively). Geneticin 600 µM or merafloxacin 50 µM were added after the infection. Cells were lysed after 24hpi or 48hpi. Protein quantification was done by Western Blot.

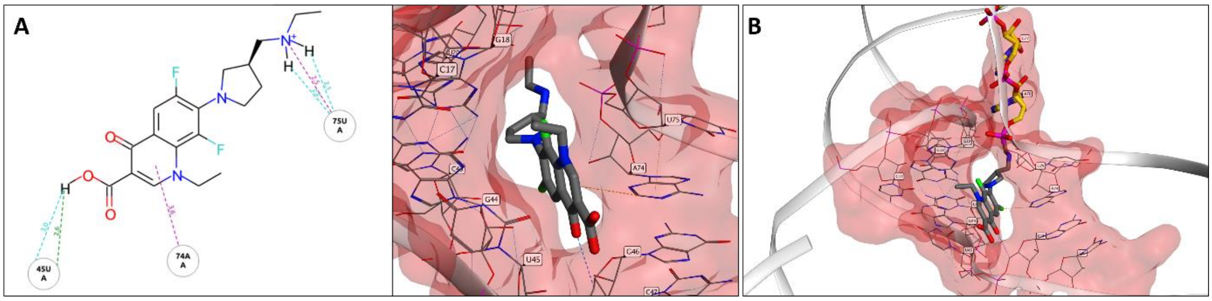


Supplementary Figure 7. A) Binding pose of merafloxicin in the site1, B) ΔACA in J3/2 region highlighted in yellow

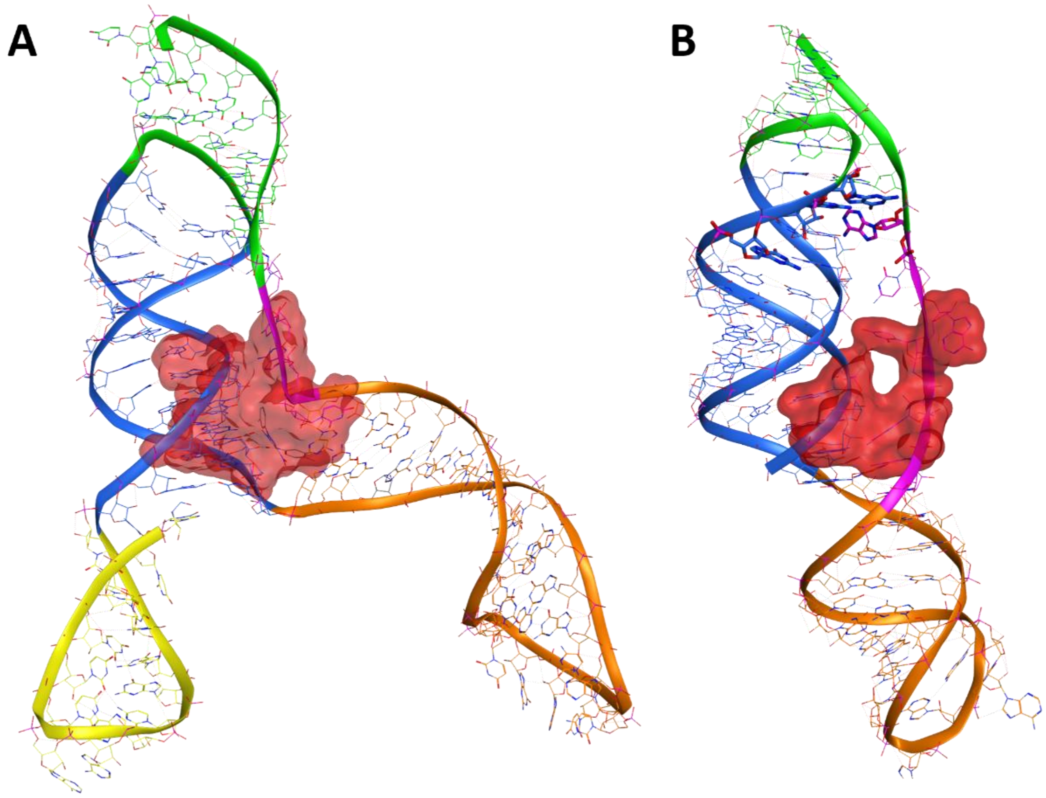


Supplementary Figure 8. Comparison between the *in silico* identified binding site in the cryo-EM structure of FSE and the recently resolved x-ray structure. In both structures, the nucleotide residues, G18, G19, G20, G43, G44, G46, U75 and A76 can form a similar binding site.

| Binding site | MM-GBSA (maestro) | Dock6 | Annapurna |
|---|---|---|---|
| 1 | -102.91 | -83.69 | -328.55 |
| 2 | -80.77 | -66.08 | -78.93 |
| 3 | -90.34 | -78.47 | -264.6 |

Supplementary Table 1. Docking score of geneticin in the three different binding sites
